# The effect of laboratory diet and feeding on growth parameters in zebrafish

**DOI:** 10.1101/2024.03.20.585913

**Authors:** Courtney Hillman, Austin H. Cooper, Pooja Ram, Matthew O. Parker

## Abstract

Despite being one of the most used laboratory species in biomedical, behavioural and physiological research, the nutritional requirements of zebrafish *(Danio rerio)* are poorly understood, and no standardised laboratory diet exists. Diet and feeding regimen can significantly impact the welfare of the fish and in turn experimental reproducibility. Consequently, the establishment of a standardised diet and feeding protocol for laboratory zebrafish is imperative to enhance animal welfare, guarantee research reproducibility and advance the economic and environmental sustainability of laboratory dietary practices. The aim of this systematic review was to determine the optimal feed for juvenile zebrafish growth and development. A comprehensive search was conducted in PubMed, Scopus and Google Scholar to identify relevant studies published up to August 2023 and the studies were selected based on the predefined inclusion/exclusion criteria. A total of 1065 articles were identified in the databases, of which 14 were included in this review. We conducted data extraction and risk-of-bias analysis in the included studies. Statistical comparisons for specific growth rate, weight gain (%) and length gain (%) parameters were performed to determine the optimal feed for enhanced juvenile growth. We identify an insect-based diet as optimal for juvenile growth for all three growth parameters. We also identify areas of potential heterogeneity and conclude by encouraging a standardised laboratory diet to ensure reproducible data and encourage zebrafish welfare.

## 1. Introduction

Zebrafish *(Danio rerio)* are one of the most common research organisms in the fields of behaviour, genetics, physiology and biomedical science with high genetic homology to mammals, low husbandry costs, rapid breeding and an ease of genetic manipulation ^1–3^. However, despite their increasing use in research, there is a lack of standardised feeding regimen and knowledge on nutritional requirements ^4–7^. The lack of consistent diet used between research groups can significantly affect fish welfare and increase experimental inconsistencies ^4,8^. It is therefore essential that a standardised diet is established which promotes welfare of the zebrafish with the added benefit of improving environmental and economic sustainability of feeding.

A lack of comprehension regarding the requirements of the optimal zebrafish diet has limited the ability to determine a standardised laboratory diet. Currently, the dietary requirements of laboratory zebrafish are related to published information on other species of fish that demonstrate similar feeding habits or live in a comparable habitat to wild zebrafish ^4,9–11^. Fig. 1 provides a basic overview of the known dietary requirements of zebrafish. It is widely accepted that dietary protein is the principal constituent for growth and development of zebrafish due to its deposition as lean body mass tissue which contributes to overall weight gain. The lipid requirements of zebrafish are unknown; however, it is assumed that they require similar amounts to other fish species. On average current diets for zebrafish constitute anywhere between 10 – 18% lipid content of dry matter ^4,12^. Carbohydrates are a principal energy source for zebrafish and therefore are considered an essential constituent of the diet. In addition, zebrafish require a range of vitamins and minerals for a balanced diet including calcium, sodium, iron, riboflavin and biotin, although the exact requirements remain to be elucidated ^4^. Like other species, any deficiencies in vitamins and minerals can result in metabolic and structural abnormalities. Although it is agreed that zebrafish require these dietary constituents for a balanced and effective diet, the overall diet composition remains unknown.

**Fig. 1.**
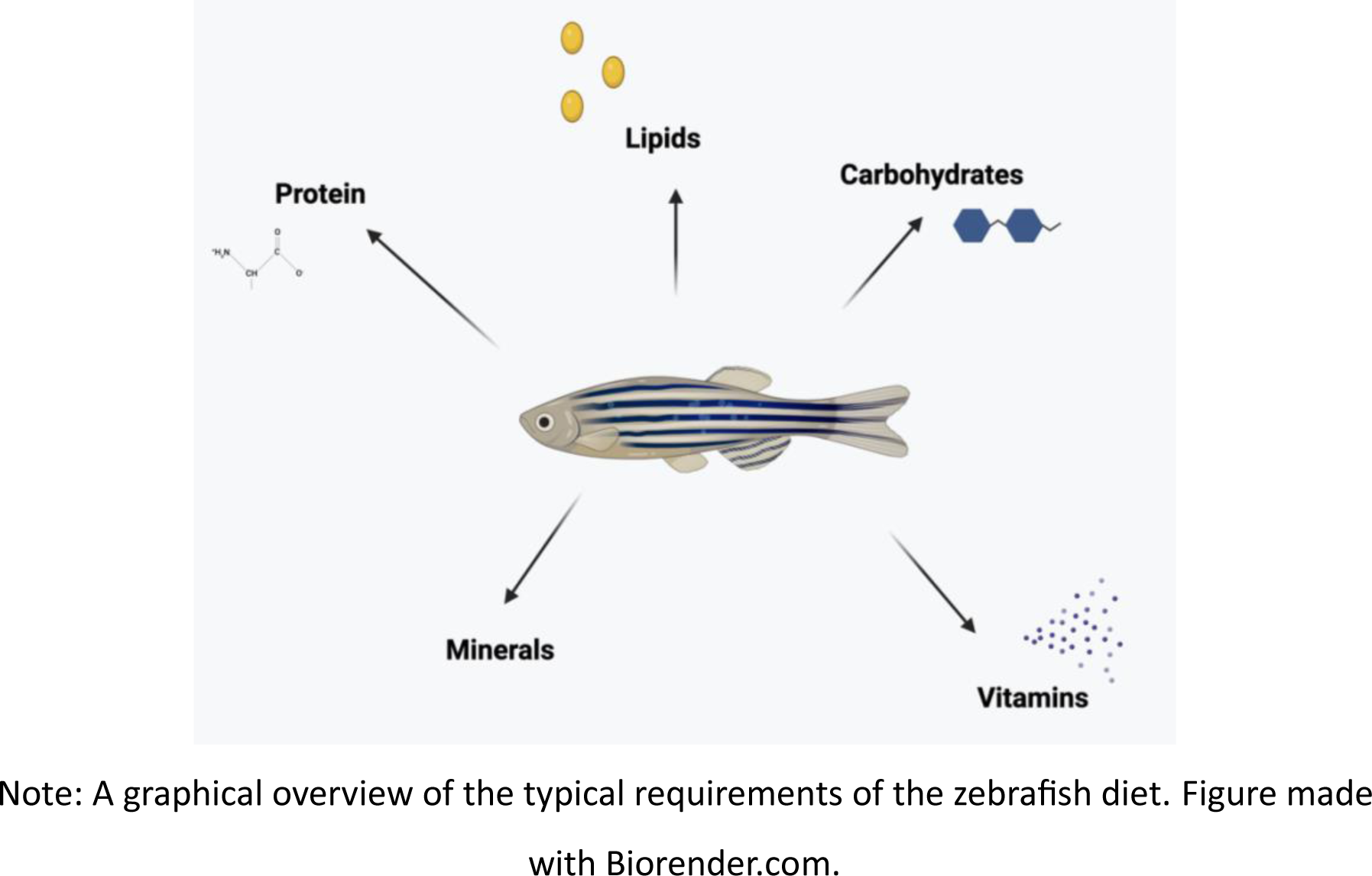
The typical constituents of a zebrafish diet.

Therefore, it is essential to determine a standardised laboratory diet through establishing the nutritional requirements of zebrafish to ensure reproducible research and promote welfare. In this review we aimed to establish the optimal feed for three different parameters of growth (specific growth rate (SGR), length gain and weight gain) in juvenile zebrafish. To determine this, we performed a systematic review of the available scientific literature studying diet on growth in juvenile zebrafish. We provide an overview of the current literature relating to zebrafish growth responses with different feeds by providing a qualitative description of the published studies as well as evaluating the impact of bias arising from methodological conduct, reporting quality and selective publication. In summary, our aim was to provide a comprehensive and data-driven approach to investigate the following research question: What feed-type is optimal for growth in juvenile zebrafish?

## 2. Methods

This systematic review was conducted in accordance with the Preferred Reporting Items for Systematic Reviews and Meta-Analyses (PRISMA) guidelines ^13^. All data and our analyses are accessible and downloadable on the Open Science Framework (OSF) (https://osf.io/d5f7t/?view_only=8445745e4716467a9997bc62700ee67f).

### 2.1. Search Strategy

Searches were carried out in three bibliographic databases -- PubMed, Scopus and Google Scholar -- using keywords which relate to our research topic for the intervention (fish feeds) and the desired population (laboratory zebrafish). Therefore, the following search string was applied: ‘Zebrafish AND (‘feed’ OR ‘diet’ OR ‘feeding’ OR ‘food’) AND (‘growth’ OR ‘survival’ OR ‘reproduction’ OR ‘welfare’). The search was carried out with no limitations on language or start date until August 2023. The reference lists of the included studies were also screened to detect additional relevant articles.

### 2.2. Eligibility screening

After searching the databases, the selection of studies included in this review was performed by one independent researcher (C.H.) with regular discussions with a second reviewer (M.P.). Titles/abstracts were initially screened to identify and exclude duplicates. Thereafter, studies were selected based on the inclusion and exclusion criteria (see below) by reading the titles/abstracts and, where necessary, the full text. Several studies had restricted access (paywalls), and the corresponding authors were contacted by email to request copies of the paper. They were given 14-days to provide the full-text of the article, after which the study was excluded from the analysis.

Studies were included if they met the following criteria: (1) original experimental research performed with adult zebrafish ≥ 28 dpf testing different fish feeds or feeding regimens; (2) studies reporting growth (SGR, length and/or weight) and/or reproductive effects of fish feeds; (3) articles were peer-reviewed; (4) fish ≥ 28 dpf but ≤ 4 months old at the start of the feeding trial (growth analysis); (5) fish > 4 months at the start of the feeding trial (reproductive analysis). Exclusion criteria were: (1) use of zebrafish < 28 dpf or other research organisms; (2) studies which incorporated over- or under-feeding; (3) assessment of food supplements for translation into higher organisms; (4) review articles, retracted articles, book chapters, scientific letters, and conference abstracts.

We required a minimum of five studies for both growth and reproduction analyses. However, during this stage we discovered that there were too few reproduction-based studies which fit our inclusion criteria and therefore the reproductive analysis was removed (See Section 3.1. Search Results for further information).

### 2.3. Data extraction

One investigator (C.H.) worked independently to complete the data extraction, and consultation was provided, when necessary, by an additional reviewer (M.P.). All data extraction was performed from the full text and figures and, where required, data were recalculated into the desired format (i.e., percentage length gain was calculated from initial and final length data). The following information was collected: (1) general data: title, authors, publication year, age of fish, strain of fish, sex split in the experiment, sample size per experimental group (n), tank density, diet and feeding regimen prior to experimental diet, experimental diet and feeding regimen and husbandry conditions (See Table 2); (2) growth results where applicable including: standard growth rate (SGR), percentage weight gain, percentage length gain and percentage survival rate (see Supplementary Data Table 4). Data were extracted as mean ± SE and when papers reported standard deviation (SD), this was converted into SE using an equation previously described in literature and used in previous meta-analyses ^14^.

Some studies only presented data in graphs (or not at all); therefore, authors were contacted via email to provide the additional data required for our analysis. These authors were given a 14-day period to respond, thereafter, PlotDigitizer (version 2.6.9. www.polotdigitizer.com) was used to manually estimate numbers from the graphs. Co-authorship networks were constructed using VOSviewer software version 1.6.20 (www.vosviewer.com) ^15^.

### 2.4. Risk of Bias and Reporting Quality

To evaluate the quality of included studies, a risk-of-bias assessment was conducted by three independent investigators (C.H., A.C. and P.R.) for each paper. This analysis was performed using the <u>SY</u>stematic <u>R</u>eview <u>C</u>entre for <u>L</u>aboratory animal <u>E</u>xperimentation (SYRCLE) risk of bias tool for animal studies ^16^. The risk of bias was assessed based on: (1) random allocation of the animals; (2) comparable baseline groups; (3) allocation randomly concealed from researchers; (4) animals randomly housed; (5) confirmation of blinded investigators and/or caregivers; (6) animals selected at random for outcome determination; (7) description of investigator blinded during outcome assessment; (8) any incomplete data justified; (9) non-selective outcome reporting; (10) any other potential biases. The reporting was as follows: low (green), slight (light green), moderate (orange), high or unclear (red). Bias plots were created using GraphPad Prism 10 (GraphPad Prism version 10.1.2 for Windows, GraphPad Software. Boston, Massachusetts USA, www.graphpad.com).

### 2.5. Data analysis

The studies were split into three groups (SGR, percentage weight gain and percentage length gain) depending on data availability. Many of the identified studies were designed as a ‘control’ feed vs ‘experimental’ feed(s) experiment. Because there was no consistency between the ‘control’ diets, we further grouped the feeds together into six distinctive categories: (1) flake/pellet feed; (2) plant-based protein; (3) animal-based protein; (4) insect-based protein; (5) supplements and additives; (6) combinations of feed/proteins. From this, where appropriate, a one-way Analysis of Variance (ANOVA) or unpaired student’s t-test was performed to determine the most effective feed within the category using GraphPad Prism. For studies where more than one feed was available within a category, the feed reported as the most effective within the study was used for the analysis. A final one-way ANOVA was conducted combining the optimal feeds from each category to determine the overall optimal feed category for SGR, percentage weight gain and percentage length gain. Type 2 error rates were *p\*\*\*\** < 0.0001, *p*\*\*\* < 0.001, *p\*\** < 0.01, *p\** < 0.05. Fig.2 provides a detailed breakdown of this analytical process.

**Fig. 2.**
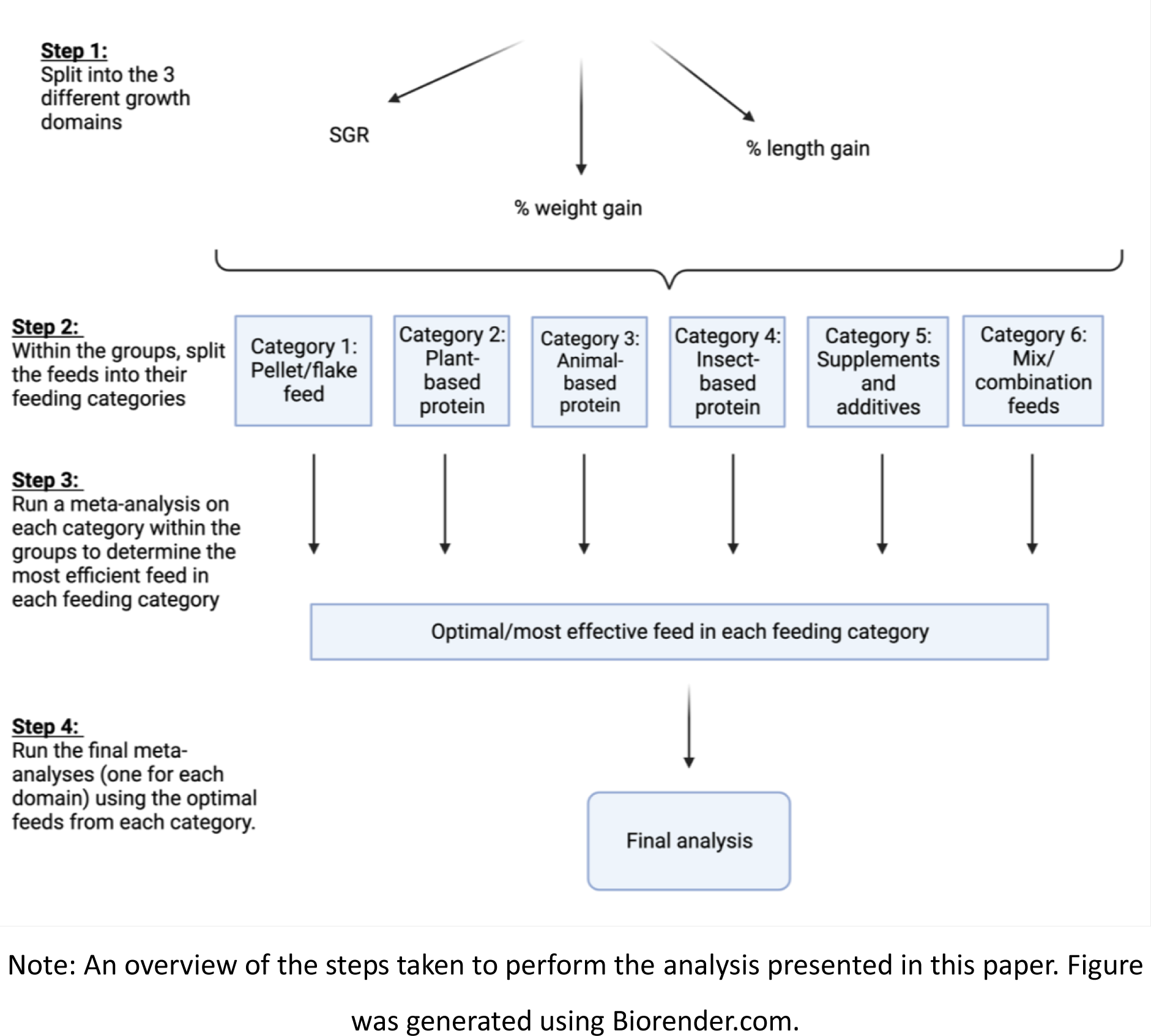
Flow chart of analytical steps use for statistical comparisons.

## 3. Results

### 3.1. Search Results

Database searches resulted in an initial 1065 documents (*n* = 512 from PubMed, and *n* = 553 from Scopus) through systematic searches (Fig. 3). Duplicates (*n* = 656) were removed and studies with unrelated research questions, different research model, and other study types were excluded (*n* = 590). After these exclusions, 66 articles remained, which were subjected to a full-text reading. Any studies which exclusively used zebrafish < 28 dpf, incorporated supplements into the diet for translational purposes or over and/or underfed the fish for obesity studies were excluded (*n* = 41). After selection by title, abstract and full text, 25 original research articles with adult zebrafish ≥ 28 dpf testing the effects of feed on growth were obtained. Fourteen of these studies were included in the present systematic review and the main characteristics are found in Table 2 and additional extracted data in supplementary data Table 4. The four studies that were not included having been initially deemed suitable were excluded because they produced significant publication bias (*n =* 1) or were the wrong age for the parameters for which they presented data (*n =* 3)^8,17–19^. Four studies were identified which assessed reproduction; however, this was below our threshold for inclusion for statistical comparison and therefore were not included any further in the analysis ^20–23^.

**Fig. 3:**
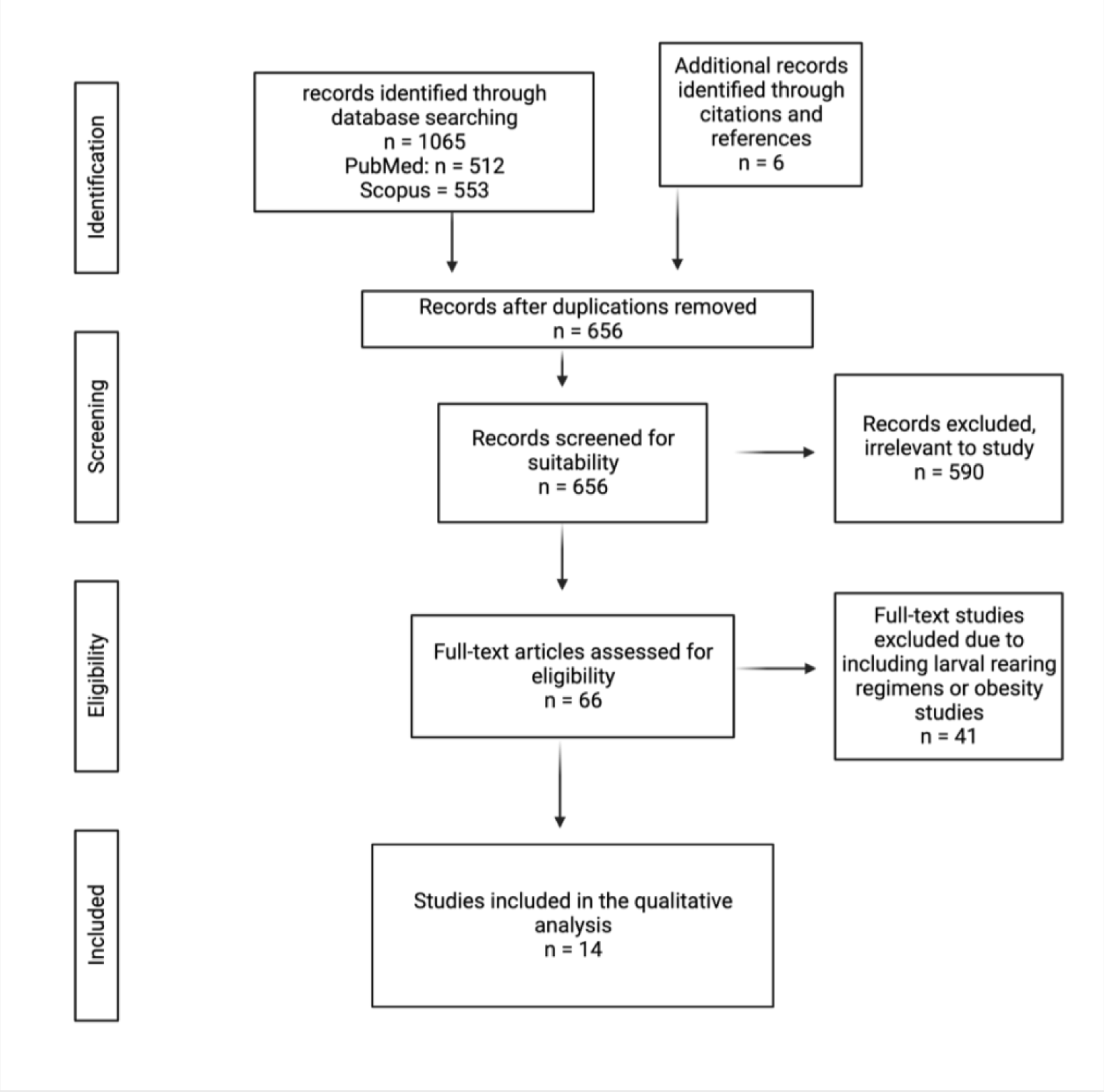
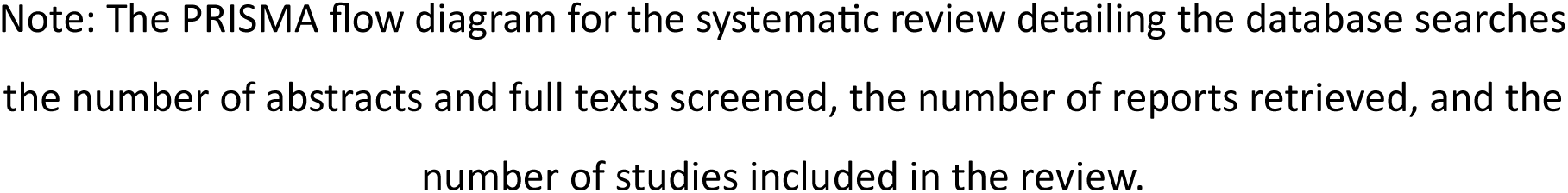
Flowchart diagram of the collection of studies and selection process.

### 3.2. Study Characteristics

The studies included in this review were published between 2012 and 2023. These studies were carried out in the United States of America (USA) (n = 3), Italy (n = 2), Turkey (n=2), India (n = 3), Brazil (n = 2), France (n = 1) and Portugal (n =1). The ages of fish ranged from 28 dpf to 4-months at the start of the feeding trial and trial length ranged from 30 to 210 days (Fig. 4). A breakdown of the diets can be seen in Table 1.

**Fig. 4:**
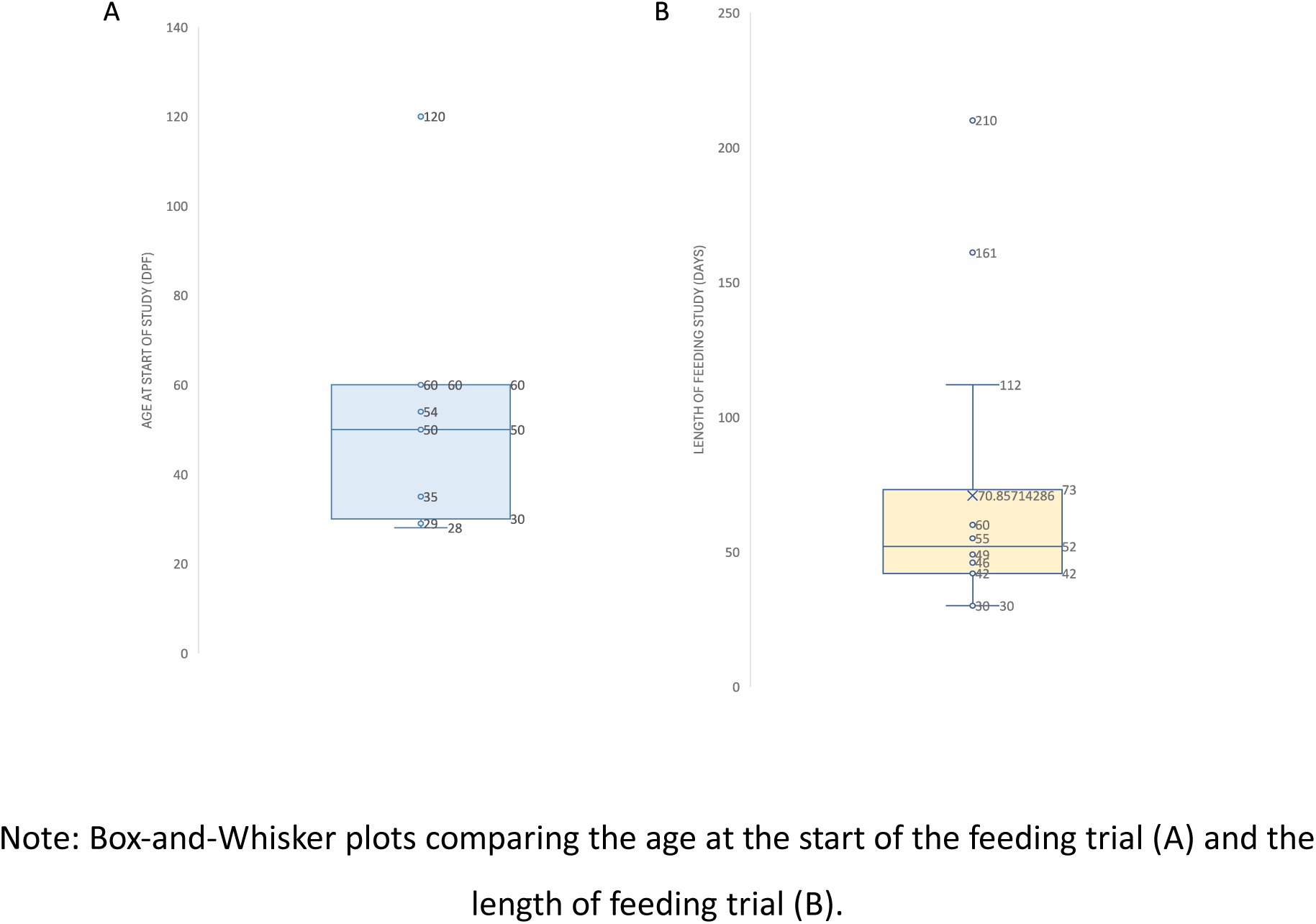
Comparison of the age at the start of the feeding trial and length of feeding trial represented using box-and-whisker plots.

**Table 1:**
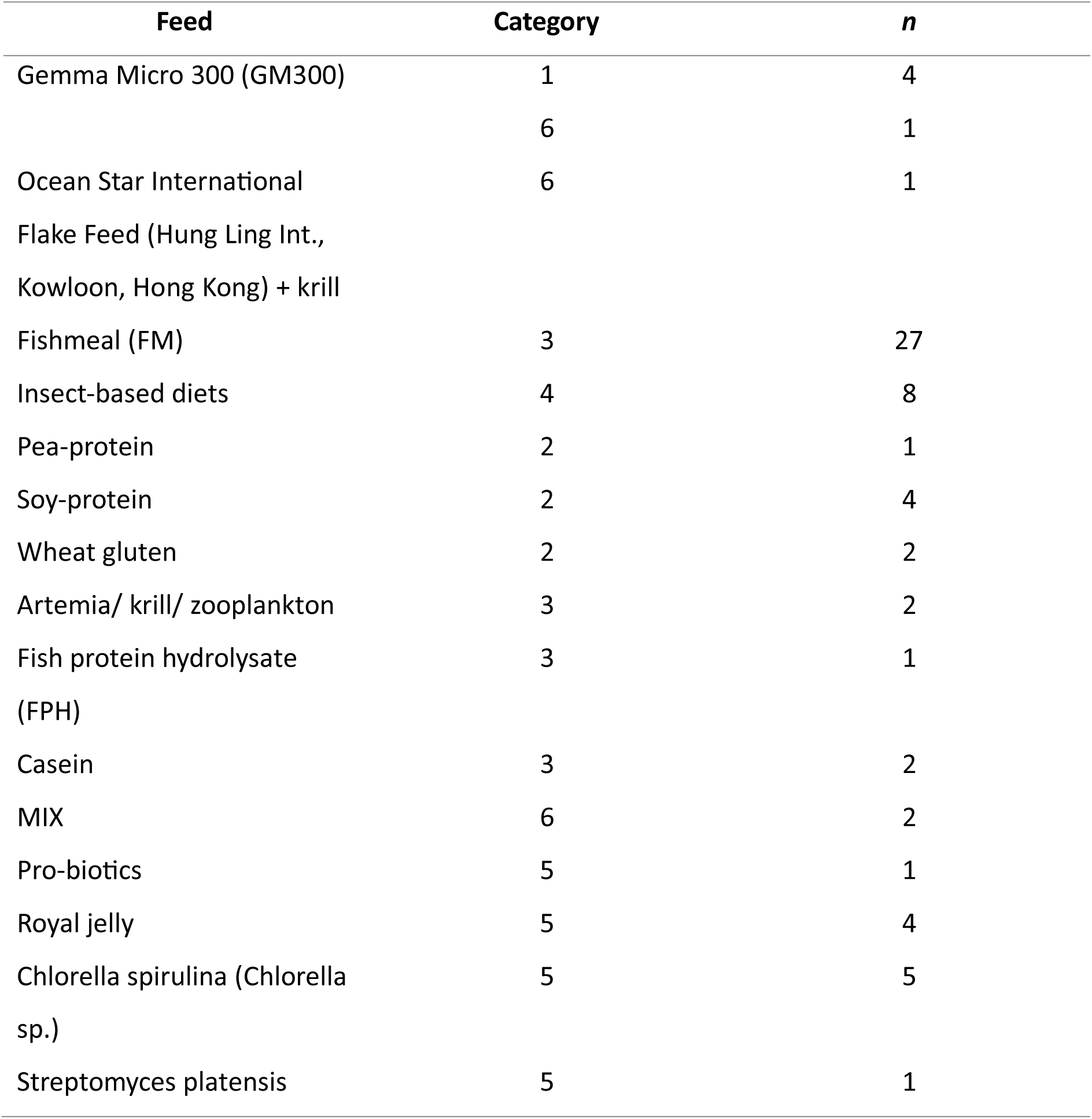
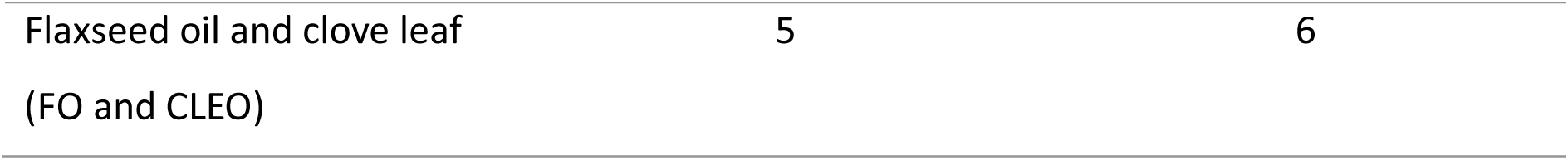
The different feeds used in the articles including the feeding category and the number of groups using this feed (*n*).

The studies reported significant effects on juvenile growth with differing zebrafish feeds with no effect on overall survival observed ^24–27^. A common finding amongst authors was a positive correlation between protein content, pro-biotic, *Chlorella spirulina (Chlorella sp.)* concentration and insect-based diets with body weight ^24,25,28,29^. Similar findings were reported on for fish length. However, feed intake was reported to decrease with increasing protein content whilst a positive relationship was observed between body weight and lean body mass with the diets ^28,30,31^. In addition to growth and survival parameters, Dhanasiri (2020) reported moderate transcriptome changes in fast-muscle samples in zebrafish fed plant-based diets compared to animal-based diets. Vural (2021) also highlighted up-regulation of growth hormone genes with Royal Jelly supplementation compared to an animal-based diet. A description of the studies included in the review can be found in Table 2. More detailed information at the study level for the variables extracted is available in Supplementary Data Table 4. The co-authorship network analysis can be found in Figure 5, showing collaboration between researchers or research groups relating to feeding effect on zebrafish growth.^32^ Limited collaboration was identified, see the interactive version of this co-authorship analysis for further information (www.vosviewer.com) ^15^.

**Table 2:** Qualitative description of studies reporting growth-related effects of experimental diets on juvenile zebrafish.

| Author, year | Age at the start of study – end of study (dpf (or months if specified)) | Sex (M:F) | Strain | Number of fish/ experimental replicate | Tank density/L | Diet(s) pre-experiment | Experimental Diet(s) | Feeding regimen | Housing conditions |
| --- | --- | --- | --- | --- | --- | --- | --- | --- | --- |
| Barca, 2023 | 60 - 102 | N/A | AB | 80 | 5.7 | N/A | D1: 50% FM<br>D2: 17% insect meal + 33% FM<br>D3: 33% insect meal + 17% FM<br>D4: 50% insect meal | *ad libitum* according to 'five-minute rule' for 42 days | Water temperature: 28 ± 0.5 °C, pH: 7.2-7.8, electrical conductivity: 600 and 800 µS/cm, dissolved oxygen: > 5 mg/L, ammonia: < 1 mg/L, nitrites: < 0.25 mg/L, nitrates: < 50 mg/L, photoperiod: 12:12 L:D, water flow: 2 L/h |
| Fredrickson, 2021 | 29 – 91 (growth at 72) | 1:1 | EK | 572 | 10.66 | 5-9 dpf: type-L saltwater rotifers<br>10-28 dpf: brine shrimp 2/day + Larval AP100 1/day + Hatchfry Encapsulon 3 1/day | D1: Ocean Star International flake food + freeze dried krill 3:1<br>D2: GM300 + brine shrimp | D1: 2/day + 2/day brine shrimp (0.2 g/10 fish)<br>D2: 1/day (10 mg/10 fish) + brine shrimp (0.2 g/10 fish) | Water temperature: 26.7 °C, pH: 7.0, electrical conductivity: 1000 µS/cm, dissolved oxygen: 7.30 mg/L, photoperiod 14:10 L:D. |
| Fronte, 2021 | 60 – 109 | N/A | AB | 60 | N/A | 4/day with commercial feed + brine shrimp until satiation following the 'five-minute rule' | D1: 20% FM<br>D2: 5% HIM + 15% FM<br>D3: 10% HIM + 10% FM<br>D4: 20% HIM | 4/day to satiation for 49 days | Water temperature: 28 ± 0.5 °C, electrical conductivity: 600-800 µS/cm, dissolved oxygen: > 7 mg/L, ammonia: 1 mg/L, nitrite: 0.25 mg/L, nitrate: 50 mg/L, photoperiod: 12:12 L:D |
| Lanes, 2021 | 30 - 60 | N/A | External supplier | 100 | 6.25 | N/A | D1: 50% FM<br>D2: defatted V instar larvae meal<br>D3: defatted prepupae meal | 3/day *ad libitum* for 30 days | Water temperature: 27 ± 0.3 °C, pH: 8.1 ± 0.1, dissolved oxygen: 6.4 ± 0.12 mg/L, photoperiod: 12:12 L:D. |
| Samuel, 2021 | Uniform sized adult | N/A | PET | 10-12 | N/A | N/A | D1: Soybean-based diet<br>D2: FM<br>D3: pro-biotic based diet<br>D4: combination<br>D5: crude protein | 1/day 4% of body weight for 30 days | N/A |
| Vural, 2021 | 2-months | 1:1 | PET | 50 | N/A | N/A | D1: casein<br>D2: 0.10% RJ<br>D3: 0.40% RJ<br>D4: 1.60% RJ<br>D5: 6.40% RJ | 4/day at 5% body weight for 56 days | Water temperature: 26 ± 1 °C, pH: 7.3 ± 0.3, dissolved oxygen: 9 ± 0.5 mg/L, photoperiod: 14:10 L:D |
| Carneiro, 2020 | 30-90 | 1:1 | short-fin | 75 | 5 | 2/ day for 2 weeks with flocculated commercial feed | D1: 50% FM<br>D2: 40% FM, 10% CS<br>D3: 30% FM, 20% CS<br>D4: 20% FM, 30% CS<br>D5: 10% FM, 40% CS<br>D6: 50% CS | 3/day until satiation for 60 days | Water temperature: 27.8 ± 0.6 °C, pH: 7.4 ± 0.3, dissolved oxygen: 7.9 ± 0.5 mg/L, ammonia: 0.09 ± 0.05 mg/L, photoperiod: 14:10 L:D |
| da Silva, 2019 | 50 - 105 | 1:0 | N/A | 60 | 1.2 | N/A | D1: Soy protein isolate<br>D2: 3% FO + 0.5% CLEO<br>D3: 3% FO + 1% CLEO<br>D4: 6% FO + 0.5% CLEO<br>D5: 6% FO + 1% CLEO<br>D6: 9% FO + 0.5% CLEO<br>D7: 9% FO + 1% CLEO | 4/day until satiation for 55 days | Water temperature: 26.23 ± 0.54°C, pH: 7.3 ± 0.01, dissolved oxygen: 6.6 ± 0.32 mg/L |
| Dhanasiri, 2020 | 4-months | 1:1 | AB | 32 | 4.57 | Live feed first + commercial feed | D1: 79.4% FM | 2/day 2.5% (w/w) of BW for 46 days | Water temperature: 28 ± 0.5°C, pH 7.5, electrical conductivity: |
| | | | | | | | D2: pea protein concentrate + FM<br>D3: WG + FM<br>D4: SPC + FM | | 1500 μS/cm, photoperiod: 12:12 L:D |
| Sevgili, 2019 | 35 | N/A | pink type | 90 | 3 | acclimated diet of commercial rainbow trout | D1: 19.87 FM<br>D2: 26.49 FM<br>D3: 30.90 FM<br>D4: 37.51 FM<br>D5: 41.92 FM<br>D5: 48.54 FM<br>D6: 52.95 FM<br>D7: 59.56 FM | *ad libitum* 2/day for 4-6 weeks | Water temperature: 24.87 ± 0.49°C, pH: 8.52 ± 0.06, dissolved oxygen: 7.65 ± 0.06 mg/L, ammonia: <0.02 mg/L, nitrites: 0.013 ± 0.003 mg/L, photoperiod: 13-14:11-10 L:D. |
| Fernandes, 2016 | 54 | N/A | PET | 20 | 18 | Brine shrimp up to 30 dpf + TetraMin flake feed 30 - 54 dpf | D2: 15% FM<br>D2: 20% FM<br>D3: 25% FM<br>D4: 30% FM<br>D5: 35% FM<br>D6: 40% FM<br>D7: 45% FM<br>D8: 50% FM<br>D9: 55% FM<br>D10: 60% FM | fed to satiation 2/day 6days/week for 8 weeks | Water temperature: 28°C ± 1°C, pH: 8.2, photoperiod: 14:10 L:D |
| Karga & Mandal, 2016 | 2-months | 1:1 | N/A | 60 | 2.5 | N/A | D1: zooplankton<br>D2: 35% FM<br>D3: 40% FM<br>D4: 45% FM | 4% BW for 90 days then 3% BW for 210 days | Water temperature: 27–28 °C, pH: 6.8 – 7.4, dissolved oxygen: 6-8 mg/L, ammonia: 0.01 – 0.6 mg/L, photoperiod: 12:12 L:D |
| Smith, 2013 | 28 | N/A | AB | 50 | 5.5 | 5-28 dpf: *Branchionus plicatilis* rotifer *ad libitum* enriched with Nannochloropsis 3/day | D1: FPH<br>D2: Casein<br>D3: SPI<br>D4: WG | 3/day at 5% BW for 16 weeks | Water temperature: 28°C, pH: 7.4, electrical conductivity: 1500 μS/cm |
|  |  |  |  |  |  |  | D5: Mix (FPH, casein, SPI, WG) |  |  |
| Lawrence,<br>2012 | 30 - 191 | 1:1 | AB | 140 | 11 | 5-9 dpf Tyle L saltwater rotifer (*Brachionus plicatilis*) *ad libitum*<br>10-30 dpf brine shrimp<br>3/day to satiation | D1: Art.<br>D2: GM300 1<br>D3: GM300 2<br>D4: GM300 3<br>D5: GM300 4 | D1: to satiation 3/day<br>D2: 5% BW 1/day<br>D3: 1.65% BW 3/day<br>D4: 1% BW 5/day<br>D5: 5% BW 1/2days<br>For 161 days | Water temperature: 26.69 ± 0.10°C, pH: 7.26 ± 0.02,<br>electrical conductivity: 1291.69 uS, dissolved oxygen: 7.9 ± 0.16 mg/L, nitrites: 0.02 ± 0.001 mg/L, nitrates: 2.79 ± 0.22 mg/L |

Note: Description of the studies included in the present meta-analysis. Key; BW: bodyweight, CS: Chlorella spirulina, CLEO: clove leaf oil, dpf: days post-fertilisation, D(1): diet(number), FM: fishmeal, FO: flaxseed oil, FPH: fish protein hydrosylate, GM300: Gemma Micro 300, HIM: Hermetia illucens meal, L:D : Light: Dark, N/A: not applicable, PET: pet-store zebrafish, RJ: Royal Jelly, SPC: soy-protein concentrate, SPI: soy-protein isolate, WG: Wheat gluten

**Fig. 5:**
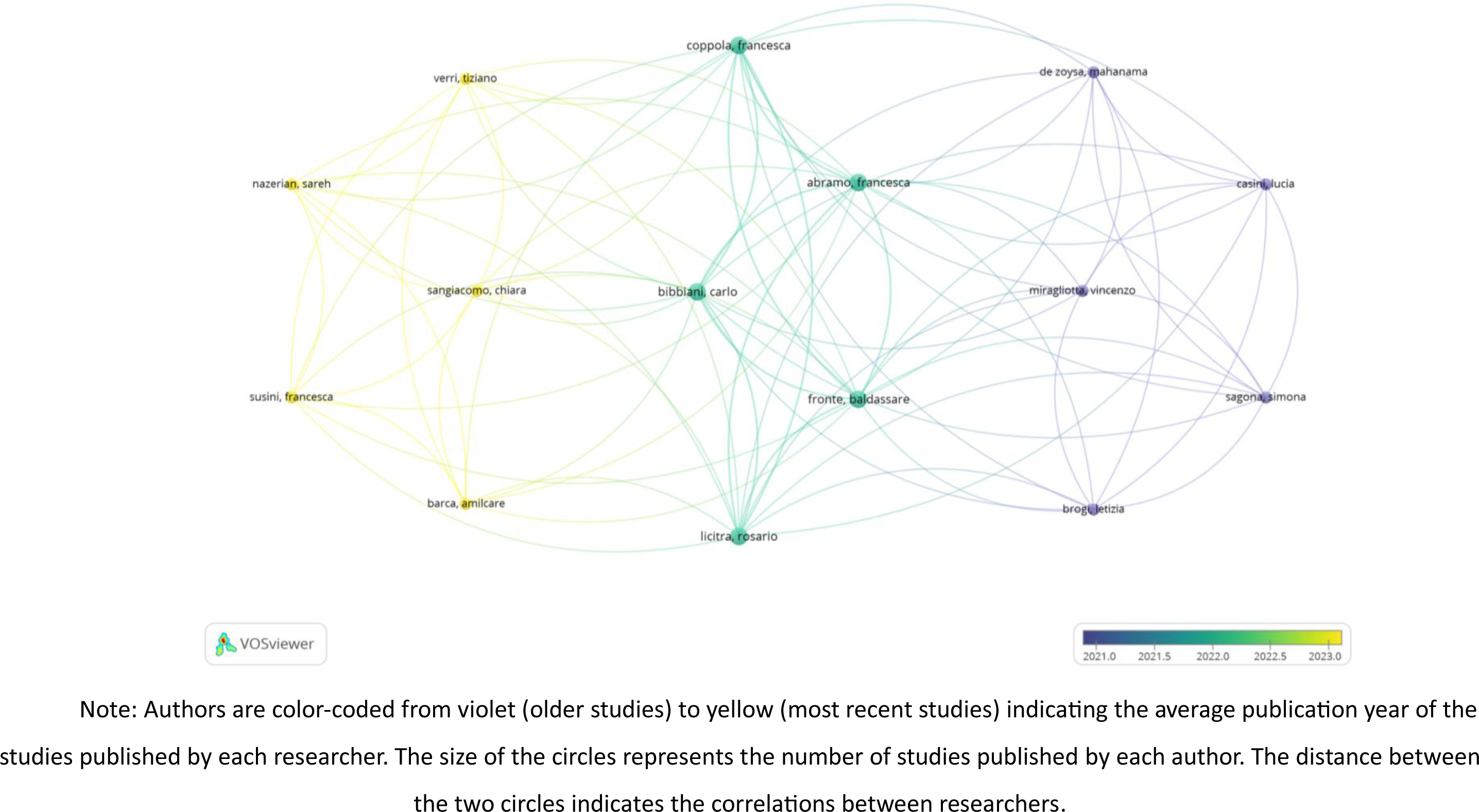
Co-authorship network analysis of researchers that authored studies assessing the effect of diet on growth.

### 3.3. Risk of bias

The overall risk of bias for the items evaluating the methodological quality of included studies was considered low, except for items 3-7, and 10 (Fig. 6). Due to this, the overall publication risk of bias presented here is considered unclear. For items 3, 5 and 7 100% of the studies provided insufficient data to rule out biases arising from the allocation of the animals to experimental groups, or biases resulting from investigators. Items 4 and 6 had moderate bias with 26.6% and 86.6% of the studies respectively providing insufficient information to rule out the potential of biases impacting results due to random housing of experimental animals and random outcome selection. Gonzales (2012) was found to have a high risk of bias for item 10 due to 50% of the experimental diets incorporating feeds which are no longer commercially available. Therefore, this study was removed from further analysis to ensure the results are of commercial relevance. Additionally, Samuel (2021) provided insufficient information regarding the age of fish with age defined as ‘uniform sized adults’ as well as a lack of husbandry condition reporting. However, this study was included in the final analysis. The eight other additional potential risks were for Barca (2023), Fronte (2021), Lanes (2021), Sevgili (2019), Fernandes (2016) and Smith (2013) who did not include a sex split and for da Silva (2020) and Karga & Mandal (2016) who did not include the strain of fish. These pose a potential risk of bias which must be taken into consideration during analysis and interpretation of results. Out of the 450 scores for risk of bias, there were 21 (4.7%) disagreements between the three independent investigators. Of the 21 disagreements, 1 (4.8%) was for item 2, 6 (28.6%) were for item 4, 5 (23.8%) were for item 6 and 9 (42.8%) were for item 10.

**Fig. 6:**
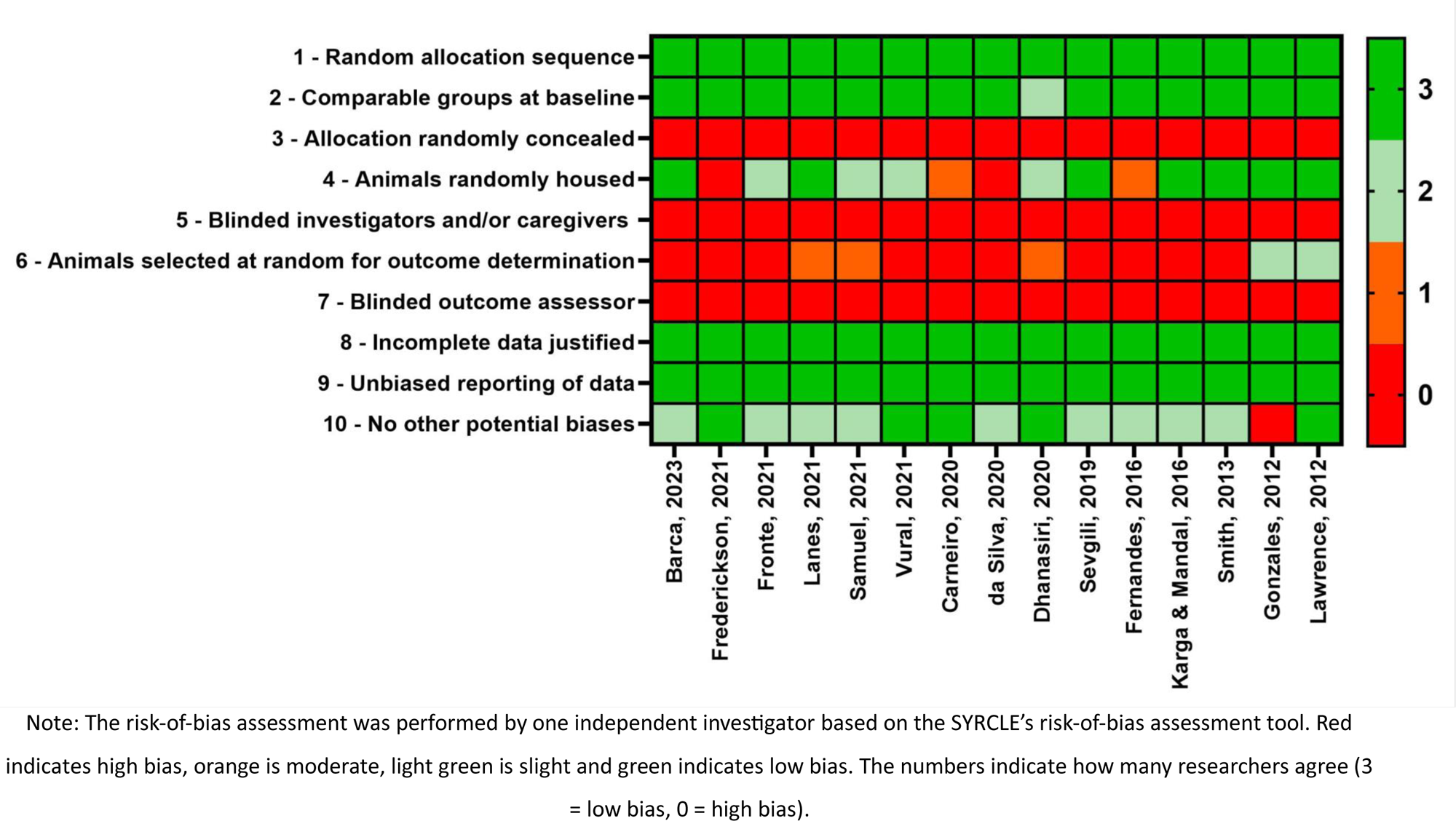
Risk-of-bias assessment of included studies.

### 3.4. Growth parameters

Zebrafish growth and development is greatly impacted by feeding regimen and protein levels; however, no standardised laboratory diet exists to date ^7,34,35^. Therefore, here we aimed to determine the optimal feed type for juvenile larval growth by performing sub-category analyses within each feeding category (Supplementary data 4-6) followed by an overall statistical analysis to determine the optimal category (Fig. 7). Optimal growth was defined here as the largest SGR, greatest percentage weight gain and percentage length gain.

**Fig. 7.**
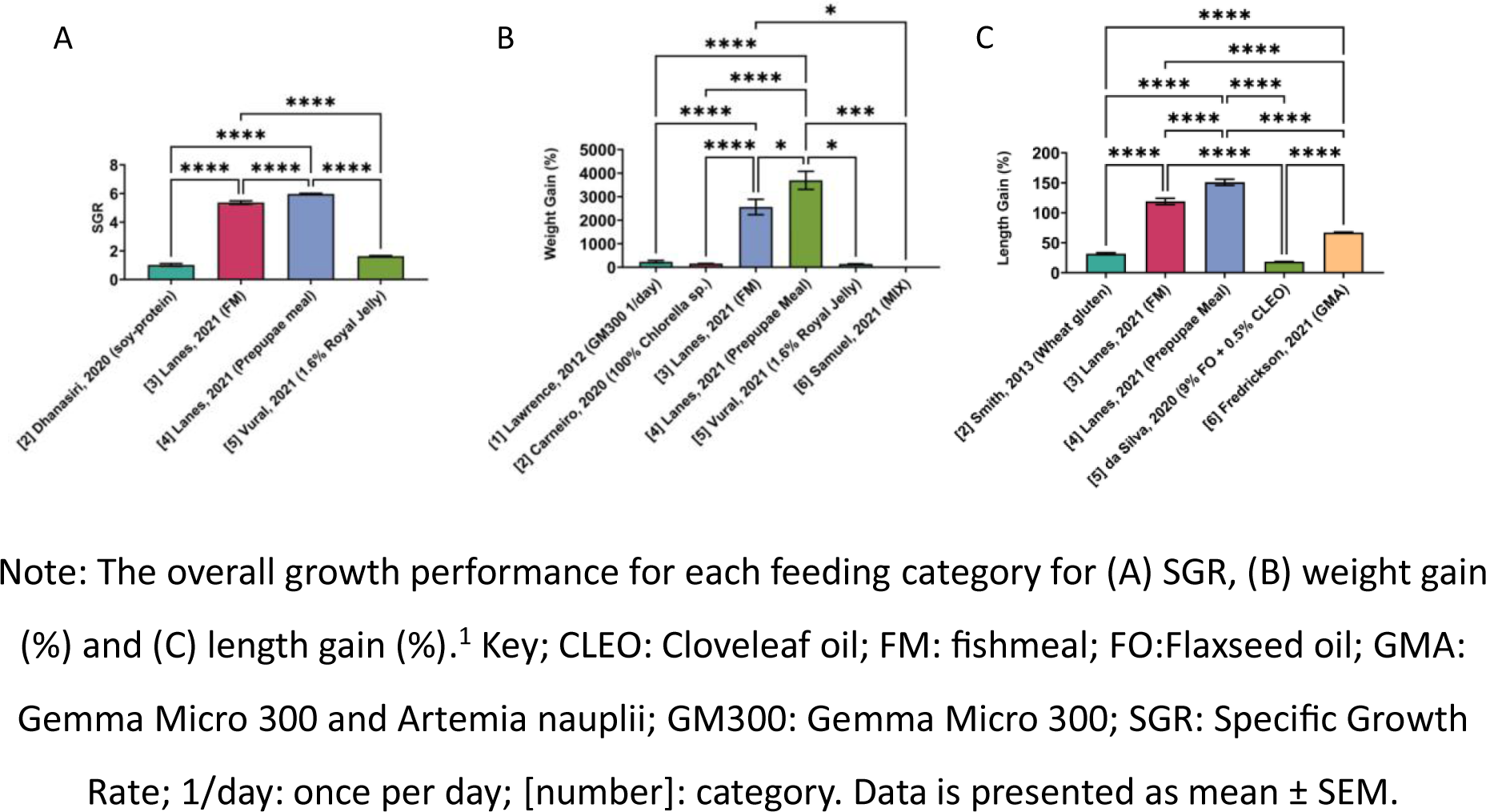
The overall growth analysis for the three growth parameters. 1 The full statistical output for all analyses can be found on the OSF for this review.

#### 3.4.1. SGR

Sub-group analysis for category 3 and category 5 SGR revealed 50% FM and 6.4% Royal Jelly to be optimal within each category, respectively (Supplementary data 4) ^36,37^. These feeds were then analysed against soy-protein isolate (category 2) and defatted prepupae meal (category 4) in the overall analysis ^37,38^. This revealed a significant effect of diet on zebrafish SGR (One-way ANOVA: F_(3, 233)_ = 311.1, *p* < 0.0001) with the optimal diet identified as category 4, defatted prepupae meal (Fig. 7A) ^37^.

#### 3.4.2. Percentage weight gain

Sub-group analysis for category 2 – 5 percentage weight gain revealed 100% Chlorella sp. (category 2), 50% FM (category 3), defatted prepupae meal (category 4) and 6.4% Royal Jelly (category 5) to be optimal within each respective category. The final analysis also included 1/day GM300 (category 1) and a protein mix (category 6) (Supplementary data 5) ^25,27,29,36,37^. The overall analysis revealed a significant effect of diet on weight gain in juvenile laboratory zebrafish (One-way ANOVA: F_(5, 424)_ = 32.09, *p* < 0.0001) (Fig. 7B). Further analysis with Tukey’s multiple comparison test revealed an insect-based diet (defatted prepupae meal) to be optimal for percentage weight gain compared to all other feeding categories, closely followed by animal-based diet with 50% FM ^37^. No significant difference was seen between the other feeding categories ^25,27,29,36^.

#### 3.4.3. Percentage length gain

Sub-group analysis revealed 50% FM (category 3) and GM300 combined with brine shrimp (category 6) to be optimal within their respective categories for juvenile length gain and were therefore compared against wheat gluten (category 2), defatted prepupae meal (category 4) and 9% FO + 0.5% CLEO (category 5) in the overall analysis ^26,30,35,37^. The analysis demonstrates a significant effect of diet on juvenile length gain (One-way ANOVA: F_(4, 877)_ = 279.6, *p* < 0.0001) (Fig. 7C). Further analysis with Tukey’s multiple comparison test revealed an insect-based diet to be optimal for percentage length gain compared to the other feeding categories (Fig. 7C). The post-hoc comparison also revealed each diet to differ significantly between each other except for wheat gluten compared to 9% FO and 0.5% CLEO ^26,30^.

## 4. Discussion

In this review, we aimed to determine the optimal feed for laboratory juvenile zebrafish growth and development. We report here that an insect-based diet, specifically defatted prepupae meal, is optimal for juvenile growth and development with significantly increased SGR, percentage weight gain and percentage length gain compared to all other tested categories (Fig. 7A-C). We also highlight the significant effect different diets can have upon growth and development of zebrafish. This signifies the importance of determining a standardised diet to promote reproducibility and experimental consistency between zebrafish laboratories.

We incorporated three different growth parameters to ensure numerous aspects of growth were accounted for within our analysis including SGR, percentage weight gain and percentage length gain. SGR is a common growth statistic used in the zebrafish community which determines the growth (unit measurement non-specific) per day ^27^. This parameter should account for the large variability in the length of feeding trial that we identified, which cannot be accounted for using weight and length gain (Fig. 4). Length and weight gain were also selected due to their commonality between the included studies. Although here we determined optimal growth to be the greatest increase in all three parameters, we do acknowledge that this may not necessarily constitute the optimal endpoint. A previous body condition scoring (BCS) has been derived for laboratory zebrafish to assess health and welfare^39^. The authors identified the optimal width of zebrafish to be between 8-10 mm ^39^. Although width was not accounted for here due to lack of reporting in studies, it highlights the importance of ensuring an optimal weight for welfare and health.

The defatted prepupae meal comes from the black soldier fly (BSF) (*Hermetia illucens* L.), which has previously been identified as a promising replacement of FM in zebrafish diet due to its comparable amino acid profile ^37^. The diet was generated from an established colony and following hatching were reared on organic waste ^37^. This highlights one of the unique environmental advantages of an insect-based diet because rearing on organic waste can reduce environmental load of waste streams ^40^. Furthermore, studies have identified the BSF for significantly reducing greenhouse gas emissions, which highlights its potential for having a substantially low carbon footprint ^41^. Additionally, incorporation of BSF in zebrafish feed could alleviate the pressures placed on fishery resources when wild fish are farmed to feed laboratory zebrafish for FM-based diets ^42^. Therefore, from an environmental perspective the incorporation of defatted prepupae meal to zebrafish diets is a suitable option.

However, the fatty acid (FA) profile of insects does not always meet the nutritional requirements of laboratory fish because insects are normally rich in saturated FA’s and mono-unsaturated FAs, rather than polyunsaturated FAs typically required by zebrafish for a balanced diet ^42^. This is a significant concern which must not be overlooked when considering the incorporation of insect-based diets for laboratory zebrafish. Insects also have chitin in their exoskeleton, which can reduce food intake, digestibility and nutrient absorption because of its limited digestive capacity ^42^. This was supported by Lanes (2021) who observed increased chitinase gene expression levels in adult zebrafish compared to their control fish fed a FM based diet. They also report that chitinase gene expression was dependent upon the BSF development stage. However, our findings presented here suggest that not only is an insect-based feed suitable for juvenile zebrafish growth and development but appears to result in substantial increases in growth compared to all other diet types. Like previous studies on zebrafish fed with BSF, we observed no significant effects on zebrafish survival (see data in OSF) ^42,43^. However, previous studies have identified that insect meal can reduce fish growth and welfare over long periods of time with a high percentage of dietary inclusion (> 25%) ^43–46^. This is suggested to be a result of the FA imbalance from the BSF ^37^. Lanes (2021) suggests that defatted BSF meal mitigates this imbalance between the FA composition of highly unsaturated and saturated FA, which is evidenced here in this review and in their research. Furthermore, a study tested defatted BSF up to 100% with Atlantic salmon (*Salmo salar*) and found comparable growth responses to control diets ^47^. Therefore, defatted BSF meal could potentially provide an economical and environmental alternative to FM with improved growth performance in juvenile zebrafish.

Although our findings suggest a significant effect of including insect-based diets for juvenile zebrafish growth and development, we acknowledge caveats to this finding. Firstly, age can have a significant effect on growth responses ^48^. Although steps were taken to reduce the likelihood of age influencing findings by providing a strict age range for the inclusion criteria, this may still have impacted the findings. The average start age between the included studies was 46 dpf (Table 2 and Fig.4). A study performed by Singleman & Holtzman (2014) suggests that zebrafish growth is greatest between the ages of 30 – 45 dpf compared to 45 – 60 dpf. However, the largest increase is seen between 90 – 180 dpf ^48^. Lanes (2021) began their study at 30 dpf and therefore may have seen a greater increase in growth compared to the average starting age of 46 dpf. The average length of feeding trial also varied significantly with Lanes (2021) having one of the shortest (Fig.4). In addition, Lanes (2021) did not account for any sex differences in growth (Table 2). Female zebrafish tend to grow heavier and longer than male zebrafish and therefore an unequal split in sex may have impacted the results ^49,50^. However, 50% of the included studies did not disclose a sex split and one of the studies that did only used males ^26^. Other factors which may have significantly impacted the growth findings include strain of fish, water quality, temperature, availability of feed and population density ^48^. All these factors differed significantly between the included studies (Table 2). This high heterogeneity between the included studies means the results presented here should be interpreted with caution. Furthermore, the main effects of the analyses were influenced by studies with a high risk of bias (Fig. 6). Although efforts were made throughout to improve the reporting quality of the research, the publication of studies adhering to measures designed to mitigate the risk of bias associated with methodological conduct is still low.

Despite the high heterogeneity and potential risk of bias between the studies reported on, we believe our findings to be of importance within the zebrafish community. Therefore, we suggest that further studies should be undertaken to identify long-term and generational effects of an insect-based diet on zebrafish. This includes genetic and behavioural assays as well as the impact of feeding on fecundity and offspring longevity.^40^ We also encourage the standardisation of zebrafish husbandry and feeding to ensure experimental reproducibility and encourage enhanced welfare.

## 5. Conclusions

In conclusion, our systematic review highlights the importance of identifying a standardised diet and feeding regimen for juvenile laboratory zebrafish growth and development. The lack of nutritional control not only raises welfare considerations but also concerns relating to the reliability and reproducibility of research outcomes. Our statistical comparison highlighted the significant variability in growth responses among studies with different feeding diets. However, our findings suggest that an insect-based diet, particularly defatted prepupae meal sourced from the BSF, holds promise for optimal growth and development of juvenile zebrafish. Despite the environmental and economic advantages of insect-based diets, challenges such as FA imbalance and chitin content must be addressed. While our review contributes valuable insights, caution is warranted due to substantial heterogeneity among studies and potential biases. Future research should focus on long-term effects, generational impacts, and comprehensive assessments of feeding strategies to ensure the welfare and scientific integrity of zebrafish research. Addressing these concerns is paramount for advancing the understanding of zebrafish biology and maximizing the model’s potential in various research fields.

## Declarations/Acknowledgments

### Ethics approval

Not applicable.

### Consent to participate

Not applicable.

### Consent to publication

Not applicable.

### Availability of data and materials

All data and analysis materials are available on the Open Science Framework (https://osf.io/d5f7t/?view_only=8445745e4716467a9997bc62700ee67f).

### Code availability

Not applicable.

### Authors contributions

Conceptualization: CH, MP. Formal analysis: CH, AC. Funding acquisition: MP. Investigation: CH, AC. Methodology: CH. Project administration: MP. Resources: MP. Supervision: MP. Visualization: CH and AC. Writing – original draft preparation: CH, AC. Writing – review & editing: CH, AC, MP.

## References

1. Stewart, A. M. et al. Molecular psychiatry of zebrafish. Mol Psychiatry 20, 2–17 (2015).

2. Choi, T.-Y., Choi, T.-I., Lee, Y.-R., Choe, S.-K. & Kim, C.-H. Zebrafish as an animal model for biomedical research. Exp Mol Med 53, 310–317 (2021).

3. Clark, K. J., Boczek, N. J. & Ekker, S. C. Stressing zebrafish for behavioral genetics. revneuro 22, 49–62 (2011).

4. Watts, S. A. & D’Abramo, L. R. Standardized reference diets for zebrafish: Addressing nutritional control in experimental methodology. Annu Rev Nutr 41, 511–527 (2021).

5. Farias, M. & Certal, A. C. Different feeds and feeding regimens have an Impact on zebrafish larval rearing and breeding performance. International Journal of Marine Biology and Research 1, 1–8 (2016).

6. Dametto, F. S. et al. Feeding regimen modulates zebrafish behavior. PeerJ 6, e5343 (2018).

7. Fowler, L. A. et al. Influence of commercial and laboratory diets on growth, body composition, and reproduction in the zebrafish *Danio rerio*. Zebrafish 16, 508–521 (2019).

8. Siccardi, A. J. et al. Growth and survival of zebrafish (*Danio rerio*) fed different commercial and laboratory diets. Zebrafish 6, 275–280 (2009).

9. Conti, F. et al. The application of synthetic flavors in zebrafish (*Danio rerio*) rearing with emphasis on attractive ones: effects on fish development, welfare, and appetite. Animals 13, 3368 (2023).

10. Lee, C. J., Paull, G. C. & Tyler, C. R. Improving zebrafish laboratory welfare and scientific research through understanding their natural history. Biological Reviews 97, 1038–1056 (2022).

11. Aleström, P. et al. Zebrafish: Housing and husbandry recommendations. Lab Anim 54, 213–224 (2020).

12. National Research Council. Nutrient Requirements of Fish and Shrimp. (National Academic Press, Washington DC, 2011).

13. Moher, D., Liberati, A., Tetzlaff, J. & Altman, D. G. Preferred reporting items for systematic reviews and meta-analyses: the PRISMA statement. BMJ 339, b2535– b2535 (2009).

14. Vesterinen, H. M. et al. Meta-analysis of data from animal studies: A practical guide. J Neurosci Methods 221, 92–102 (2014).

15. Kirby, A. Exploratory bibliometrics: Using VOSviewer as a preliminary research tool. Publications 11, 10 (2023).

16. Hooijmans, C. R. et al. SYRCLE’s risk of bias tool for animal studies. BMC Med Res Methodol 14, 43 (2014).

17. Gonzales, J. M. Preliminary evaluation on the effects of feeds on the growth and early reproductive performance of zebrafish (*Danio rerio*). J Am Assoc Lab Anim Sci 51, 412– 7 (2012).

18. Nowosad, J., Kucharczyk, D. & Targońska, K. Enrichment of zebrafish *Danio rerio* (Hamilton, 1822) diet with polyunsaturated fatty acids improves fecundity and larvae quality. Zebrafish 14, 364–370 (2017).

19. Gonzales, J. M. & Law, S. H. W. Feed and feeding regime affect growth rate and gonadosomatic index of adult zebrafish (*Danio rerio*). Zebrafish 10, 532–40 (2013).

20. Tye, M. et al. Nonhatching decapsulated *artemia* cysts as a replacement to *artemia* nauplii in juvenile and adult zebrafish culture. Zebrafish 12, 457–461 (2015).

21. Geffroy, B. & Simon, O. Effects of a Spirulina platensis-based diet on zebrafish female reproductive performance and larval survival rate. Cybium 37, 31–38 (2013).

22. Şişman, T., et al. Single-cell protein as an alternative food for zebrafish, *Danio rerio*: a toxicological assessment. Toxicol Ind Health 29, 792–799 (2013).

23. Markovich, M. L., Rizzuto, N. V. & Brown, P. B. Diet affects spawning in zebrafish. Zebrafish 4, 69–74 (2007).

24. Barca, A., et al. *Hermetia illucens* for replacing fishmeal in aquafeeds: Effects on fish growth performance, intestinal morphology, and gene expression in the zebrafish (*Danio rerio*) model. Fishes 8, 127 (2023).

25. Carneiro, W. F. et al. Replacing fish meal by Chlorella sp. meal: Effects on zebrafish growth, reproductive performance, biochemical parameters and digestive enzymes. Aquaculture 528, 735612 (2020).

26. da Silva, T. C., et al. Flaxseed oil and clove leaf essential oil in Zebrafish diet (Danio rerio). Acta Sci 43, e48126 (2020).

27. Lawrence, C., Best, J., James, A. & Maloney, K. The effects of feeding frequency on growth and reproduction in zebrafish (*Danio rerio*). Aquaculture 368–369, 103–108 (2012).

28. Fernandes, H., Peres, H. & Carvalho, A. P. Dietary protein requirement during juvenile growth of zebrafish (*Danio rerio*). Zebrafish 13, 548–555 (2016).

29. Samuel, R. A., Sasmita Dash, S., Ali, L. R. & Alagesan Paari, K. Formulation and characterization of plant, animal, and probiotic based fish meals and evaluating their efficacy on growth and performance in zebrafish (*Danio rerio*). Adv Anim Vet Sci 9, (2021).

30. Smith Jr., D. L., et al. Dietary protein source influence on body size and composition in growing zebrafish. Zebrafish 10, 439–446 (2013).

31. Sevgİlİ, H., Sezen, S., Kanyilmaz, M., Aktaş, Ö. & Pak, F. Dietary Protein Requirements of Zebrafish (*Dania rerio*). Journal of Limnology and Freshwater Fisheries Research 5, 34–40 (2019).

32. Fonseca, B. de P. F. e, Sampaio, R. B., Fonseca, M. V. de A. & Zicker, F. Co-authorship network analysis in health research: method and potential use. Health Res Policy Syst 14, 34 (2016).

33. Fronte, B. et al. Fishmeal replacement with *Hermetia illucens* meal in aquafeeds: Effects on zebrafish growth performances, intestinal morphometry, and enzymology. Fishes 6, 28 (2021).

34. Valentine, S. & Kwasek, K. Feeding rate and protein quality differentially affect growth and feeding efficiency response variables of zebrafish (*Danio rerio*). Zebrafish 19, 94– 103 (2022).

35. Frederickson, S. C. et al. Comparison of juvenile feed protocols on growth and spawning in zebrafish. Journal of the American Association for Laboratory Animal Science 60, 298–305 (2021).

36. Vural, O., Silici, S. & Aksakal, E. The effect of royal jelly dietary on growth performance and expression of genes related to growth and immunity of zebrafish, Danio rerio. Aquac Rep 20, 100652 (2021).

37. Lanes, C. F. C. et al. Black soldier fly (*Hermetia illucens*) larvae and prepupae defatted meals in diets for zebrafish (*Danio rerio*). Animals 11, 720 (2021).

38. Dhanasiri, A. K. S. et al. Plant-based diets induce transcriptomic changes in muscle of zebrafish and atlantic salmon. Front Genet 11, (2020).

39. Clark, T. S., Pandolfo, L. M., Marshall, C. M., Mitra, A. K. & Schech, J. M. Body condition scoring for adult zebrafish (*Danio rerio*). J Am Assoc Lab Anim Sci 57, 698–702 (2018).

40. Spranghers, T. et al. Gut antimicrobial effects and nutritional value of black soldier fly (*Hermetia illucens L.)* prepupae for weaned piglets. Anim Feed Sci Technol 235, 33–42 (2018).

41. Pang, W. et al. Reducing greenhouse gas emissions and enhancing carbon and nitrogen conversion in food wastes by the black soldier fly. J Environ Manage 260, 110066 (2020).

42. Zarantoniello, M. et al. Partial dietary inclusion of *Hermetia illucens* (Black Soldier Fly) full-fat prepupae in zebrafish feed: biometric, histological, biochemical, and molecular implications. Zebrafish 15, 519–532 (2018).

43. Zarantoniello, M. et al. A six-months study on Black Soldier Fly (*Hermetia illucens*) based diets in zebrafish. Sci Rep 9, 8598 (2019).

44. Sánchez-Muros, M.-J., Barroso, F. G. & Manzano-Agugliaro, F. Insect meal as renewable source of food for animal feeding: a review. J Clean Prod 65, 16–27 (2014).

45. Gasco, L. et al. Tenebrio molitor meal in diets for European sea bass (Dicentrarchus labrax L.) juveniles: Growth performance, whole body composition and in vivo apparent digestibility. Anim Feed Sci Technol 220, 34–45 (2016).

46. Kroeckel, S. et al. When a turbot catches a fly: Evaluation of a pre-pupae meal of the Black Soldier Fly (*Hermetia illucens*) as fish meal substitute — Growth performance and chitin degradation in juvenile turbot (*Psetta maxima*). Aquaculture 364–365, 345–352 (2012).

47. Belghit, I. et al. Black soldier fly larvae meal can replace fish meal in diets of sea-water phase Atlantic salmon (*Salmo salar*). Aquaculture 503, 609–619 (2019).

48. Singleman, C. & Holtzman, N. G. Growth and maturation in the zebrafish, Danio rerio: a staging tool for teaching and research. Zebrafish 11, 396–406 (2014).

49. Eaton, R. C. & Farley, R. D. Growth and the Reduction of Depensation of Zebrafish, Brachydanio rerio, Reared in the Laboratory. Copeia 1974, 204 (1974).

50. Spence, R. & Smith, C. Mating preference of female zebrafish, Danio rerio, in relation to male dominance. Behavioral Ecology 17, 779–783 (2006).

