## Supplementary data for "The effect of laboratory diet and feeding on growth parameters in zebrafish"

7. **Supplementary data**
   7.1. **Supplementary data 1**

The PRISMA checklist for requirements of systematic reviews and meta-analysis is found in supplementary data table 1 including the location in the present review and the abstract checklist is in supplementary data table 2.

**Supplementary Data Table 1: The PRISMA checklist for systematic reviews and meta-analyses for this present review.**

| **Section and Topic** | **Item #** | **Checklist item** | **Location where item is reported** |
| --- | --- | --- | --- |
| **TITLE** | | |  |
| Title | 1 | Identify the report as a systematic review. | Title |
| **ABSTRACT** | | |  |
| Abstract | 2 | See the PRISMA 2020 for Abstracts checklist. | Supplementary data |
| **INTRODUCTION** | | |  |
| Rationale | 3 | Describe the rationale for the review in the context of existing knowledge. | 1.0 |
| Objectives | 4 | Provide an explicit statement of the objective(s) or question(s) the review addresses. | 1.0 |
| **METHODS** | | |  |
| Eligibility criteria | 5 | Specify the inclusion and exclusion criteria for the review and how studies were grouped for the syntheses. | 2.2 |
| Information sources | 6 | Specify all databases, registers, websites, organisations, reference lists and other sources searched or consulted to identify studies. Specify the date when each source was last searched or consulted. | 2.1 |
| Search strategy | 7 | Present the full search strategies for all databases, registers and websites, including any filters and limits used. | 2.1 |
| Selection process | 8 | Specify the methods used to decide whether a study met the inclusion criteria of the review, including how many reviewers screened each record and each report retrieved, whether they worked independently, and if applicable, details of automation tools used in the process. | 2.2 |
| Data collection process | 9 | Specify the methods used to collect data from reports, including how many reviewers collected data from each report, whether they worked independently, any processes for obtaining or confirming data from study investigators, and if applicable, details of automation tools used in the process. | 2.3 |
| Data items | 10a | List and define all outcomes for which data were sought. Specify whether all results that were compatible with each outcome domain in each study were sought (e.g. for all measures, time points, analyses), and if not, the methods used to decide which results to collect. | 2.3 |
|  | 10b | List and define all other variables for which data were sought (e.g. participant and intervention characteristics, funding sources). Describe any assumptions made about any missing or unclear information. | 2.3 |
| Study risk of bias assessment | 11 | Specify the methods used to assess risk of bias in the included studies, including details of the tool(s) used, how many reviewers assessed each study and whether they worked independently, and if applicable, details of automation tools used in the process. | 2.4 |
| Effect measures | 12 | Specify for each outcome the effect measure(s) (e.g. risk ratio, mean difference) used in the synthesis or presentation of results. | 2.5 |
| Synthesis methods | 13a | Describe the processes used to decide which studies were eligible for each synthesis (e.g. tabulating the study intervention characteristics and comparing against the planned groups for each synthesis (item #5)). | 2.5 |
|  | 13b | Describe any methods required to prepare the data for presentation or synthesis, such as handling of missing summary statistics, or data conversions. | 2.5 |
|  | 13c | Describe any methods used to tabulate or visually display results of individual studies and syntheses. | 2.5 |
|  | 13d | Describe any methods used to synthesize results and provide a rationale for the choice(s). If meta-analysis was performed, describe the model(s), method(s) to identify the presence and extent of statistical heterogeneity, and software package(s) used. | 2.5 |
|  | 13e | Describe any methods used to explore possible causes of heterogeneity among study results (e.g. subgroup analysis, meta-regression). | 2.5 |
|  | 13f | Describe any sensitivity analyses conducted to assess robustness of the synthesized results. | 2.5 |
| Reporting bias assessment | 14 | Describe any methods used to assess risk of bias due to missing results in a synthesis (arising from reporting biases). | 2.5 |
| Certainty assessment | 15 | Describe any methods used to assess certainty (or confidence) in the body of evidence for an outcome. | 2.5 |
| **RESULTS** | | |  |
| Study selection | 16a | Describe the results of the search and selection process, from the number of records identified in the search to the number of studies included in the review, ideally using a flow diagram. | 3.1 |
|  | 16b | Cite studies that might appear to meet the inclusion criteria, but which were excluded, and explain why they were excluded. | 3.1 |
| Study characteristics | 17 | Cite each included study and present its characteristics. | 3.2 |
| Risk of bias in studies | 18 | Present assessments of risk of bias for each included study. | 3.3 |
| Results of individual studies | 19 | For all outcomes, present, for each study: (a) summary statistics for each group (where appropriate) and (b) an effect estimate and its precision (e.g. confidence/credible interval), ideally using structured tables or plots. | Supplementary data |
| Results of syntheses | 20a | For each synthesis, briefly summarise the characteristics and risk of bias among contributing studies. | 3.4/5 |
|  | 20b | Present results of all statistical syntheses conducted. If meta-analysis was done, present for each the summary estimate and its precision (e.g. confidence/credible interval) and measures of statistical heterogeneity. If comparing groups, describe the direction of the effect. | 3.4/5 |
|  | 20c | Present results of all investigations of possible causes of heterogeneity among study results. | 5 |
|  | 20d | Present results of all sensitivity analyses conducted to assess the robustness of the synthesized results. | N/A |
| Reporting biases | 21 | Present assessments of risk of bias due to missing results (arising from reporting biases) for each synthesis assessed. | 3.3 |
| Certainty of evidence | 22 | Present assessments of certainty (or confidence) in the body of evidence for each outcome assessed. | 3.5/6 |
| **DISCUSSION** | | |  |
| Discussion | 23a | Provide a general interpretation of the results in the context of other evidence. | 4 |
|  | 23b | Discuss any limitations of the evidence included in the review. | 4 |
|  | 23c | Discuss any limitations of the review processes used. | 4 |
|  | 23d | Discuss implications of the results for practice, policy, and future research. | 4 |
| **OTHER INFORMATION** | | |  |
| Registration and protocol | 24a | Provide registration information for the review, including register name and registration number, or state that the review was not registered. | Review was not registered |
|  | 24b | Indicate where the review protocol can be accessed, or state that a protocol was not prepared. | OSF |
|  | 24c | Describe and explain any amendments to information provided at registration or in the protocol. | N/A |
| Support | 25 | Describe sources of financial or non-financial support for the review, and the role of the funders or sponsors in the review. | declarations |
| Competing interests | 26 | Declare any competing interests of review authors. | The authors declare no competing interests |
| Availability of data, code and other materials | 27 | Report which of the following are publicly available and where they can be found: template data collection forms; data extracted from included studies; data used for all analyses; analytic code; any other materials used in the review. | OSF |

**Supplementary Data Table 2: The PRISMA abstract checklist for systematic reviews and meta-analyses for the present review.**

| **Section and Topic** | **Item #** | **Checklist item** | **Reported (Yes/No)** |
| --- | --- | --- | --- |
| **TITLE** | | |  |
| Title | 1 | Identify the report as a systematic review. | Yes |
| **BACKGROUND** | | |  |
| Objectives | 2 | Provide an explicit statement of the main objective(s) or question(s) the review addresses. | Yes |
| **METHODS** | | |  |
| Eligibility criteria | 3 | Specify the inclusion and exclusion criteria for the review. | Yes |
| Information sources | 4 | Specify the information sources (e.g. databases, registers) used to identify studies and the date when each was last searched. | Yes |
| Risk of bias | 5 | Specify the methods used to assess risk of bias in the included studies. | Yes |
| Synthesis of results | 6 | Specify the methods used to present and synthesise results. | Yes |
| **RESULTS** | | |  |
| Included studies | 7 | Give the total number of included studies and participants and summarise relevant characteristics of studies. | Yes |
| Synthesis of results | 8 | Present results for main outcomes, preferably indicating the number of included studies and participants for each. If meta-analysis was done, report the summary estimate and confidence/credible interval. If comparing groups, indicate the direction of the effect (i.e. which group is favoured). | Yes |
| **DISCUSSION** | | |  |
| Limitations of evidence | 9 | Provide a brief summary of the limitations of the evidence included in the review (e.g. study risk of bias, inconsistency and imprecision). | Yes |
| Interpretation | 10 | Provide a general interpretation of the results and important implications. | Yes |
| **OTHER** | | |  |
| Funding | 11 | Specify the primary source of funding for the review. | (declarations) |
| Registration | 12 | Provide the register name and registration number. | N/A |

7.2. **Supplementary data 2**

The table including the searches conducted number of hits and number of articles found that are included in the analysis is reported on in Supplementary Data Table 3.

**Supplementary Data Table 3: The searches conducted, number of hits and the number of articles found which were included in the present analysis.**

| **Date of Search** | **Search String** | **Database** | **Number of hits** | **Number of articles included in analysis** |
| --- | --- | --- | --- | --- |
| 18/08/2023 | Zebrafish AND diet AND growth | PubMed | 285 | 6 |
| 21/08/2023 | Zebrafish AND diet AND welfare | PubMed | 15 | 0 |
| 21/08/2023 | Zebrafish AND diet AND health | PubMed | 212 | 0 |
| 21/08/2023 | Zebrafish AND feed AND welfare | PubMed | 14 | 0 |
| 21/08/2023 | Zebrafish OR 'danio rerio' AND feed OR diet OR feeding AND welfare | Scopus | 14 | 0 |
| 21/08/2023 | zebrafish AND feed OR diet AND reproduction OR survival OR growth | Scopus | 525 | 7 |
| 22/08/2023 | N/A | citations and references | N/A | 2 |

7.3. **Supplementary data 3**

The statistical data that was used in the analysis included in this review is seen in Supplementary Data Table 4.

**Supplementary Data Table 4: The experimental data used in the current analysis from the studies describing the growth and survival effects of different feeds on juvenile zebrafish.**

| Author, year | SGR (% ± SE) | Weight gain (% ± SE) | Standard length gain (% ± SE) | Survival rate (% ± SE) |
| --- | --- | --- | --- | --- |
| Barca, 2023^*^ |  | D1: 360.78 ± 7.68  D2: 392.25 ± 6.86  D3: 392.52 ± 7.09  D4: 417.91 ± 7.14 |  | D1: 95  D2: 98.5  D3: 100  D4: 93.6 |
| Fronte, 2021 |  | D1: 169.64 ± 9.98  D2: 178.94 ± 10.44  D3: 181.63 ± 10.08  D4: 166.16 ± 9.74 |  | D1: 88.75  D2: 92.5  D3: 88.75  D4: 93.75 |
| Lanes, 2021 | D1: 5.37 ± 0.12  D2: 5.67 ± 0.14  D3: 5.97 ± 0.04 | D1: 2559.8 ± 20.81  D2: 3088.2 ± 26.45  D3:  3694 ± 10.07 | D1: 10.94 ± 0.6986  D2: 12.64 ± 0.7431  D3: 13.88 ± 0.3970 | D1: 92.50 ± 3.23  D2: 93.75 ± 4.73  D3: 96.25 ± 2.39 |
| Samuel, 2021 |  | D1: 14.38 ± 0.2  D2: 14.0 ± 0.01  D3: 15.7 ± 0.08  D4: 14.99 ± 0.11  D5: 13.27 ± 0.15 |  |  |
| Vural, 2021 | D1: 1.49 ± 0.05  D2: 1.54 ± 0.09  D3: 1.07 ± 0.09  D4: 1.63 ± 0.04  D5: 1.54 ± 0.03 | D1: 230.14 ± 6.90  D2: 237.04 ± 11.85  D3: 182.04 ± 8.80  D4: 249.11 ± 5.45  D5: 237.01 ± 4.12 |  | D1: 79.33 ± 6.77  D2: 82.67 ± 5.81  D3: 88.0 ± 1.15  D4: 78.67 ± 10.48  D5: 84.00 ± 1.15 |
| Carneiro, 2020 | D1: 1.55 ± 0.0  D2: 1.61 ± 0.0  D3: 1.56 ± 0.0  D4: 1.55 ± 0.0  D5: 1.66 ± 0.0  D6: 1.58 ± 0.0 | D1: 153.12 ± 2.3  D2: 163.87 ± 7.7  D3: 155.56 ± 2.5  D4: 154.27 ± 4.6  D5: 170.02 ± 2.9  D6: 158.39 ± 4.2 |  | D1: 88.6 ± 3.6  D2: 94.3 ± 4.2  D3: 88.6 ± 2.9  D4: 88.6 ± 2.9  D5: 87.1 ± 3.5  D6: 91.5 ± 2.7 |
| da Silva, 2020 | D1: 0.46 ± 0.07  D2: 0.74 ± 0.15  D3: 0.78 ± 0.14  D4: 1.15 ± 0.15  D5: 0.85 ± 0.09  D6: 1.04 ± 0.13  D7: 0.93 ± 0.12 |  | D1: 31.07 ± 1.65  D2: 33.23 ± 1.29  D3: 34.16 ± 1.59  D4: 34.79 ± 1.12  D5: 34.23 ± 1.29  D6: 36.38 ± 1.35  D7: 36.89 ± 1.02 | D1: 100  D2: 100  D3: 100  D4: 100  D5: 100  D6: 100  D7: 100 |
| Dhanasari, 2020 | D1: 1.0679 ± 0.085  D2: 0.9119 ± 0.045  D3: 1.0157 ± 0.097  D4: 1.0756 ± 0.139 |  |  |  |
| Sevgili, 2018 | D1: 2.90 ± 0.075  D2: 2.91 ± 0.075  D3: 3.05 ± 0.075  D4: 3.00 ± 0.075  D5: 3.02 ± 0.075  D6: 2.83 ± 0.075  D7: 2.96 ± 0.075  D8: 2.88 ± 0.075 | D1: 238.22 ± 9.43  D2: 239.66 ± 9.47  D3: 260.00 ± 10.0  D4: 252.40 ± 10.10  D5: 254.81 ± 10.20  D6: 228.46 ± 10.17  D7: 247.05 ± 10.49  D8: 234.78 ± 10.35 |  |  |
| Fernandes, 2016 | D1: 2.4 ± 0.08  D2: 2.6 ± 0.08  D3: 2.8 ± 0.08  D4: 2.9 ± 0.08  D5: 3.2 ± 0.08  D6: 3.2 ± 0.08  D7: 3.4 ± 0.08  D8: 3.3 ± 0.08  D9: 3.4 ± 0.08  D10: 3.3 ± 0.08 | D1: 37.43 ± 2.85  D2: 40.00 ± 2.83  D3: 42.54 ± 2.93  D4: 43.55 ± 2.97  D5: 46.54 ± 2.88  D6: 47.34 ± 2.99  D7: 47.78 ± 3.00  D8: 46.79 ± 3.00  D9: 48.00 ± 3.03  D10: 47.29 ± 3.03 | D1: 44.57 ± 0.705  D2: 54.35 ± 0.800  D3: 56.52 ± 0.826  D4: 58.70 ± 0.846  D5: 66.85 ± 0.920  D6: 66.30 ± 0.906  D7: 69.57 ± 0.937  D8: 73.37 ± 0.963  D9: 72.83 ± 0.958  D10: 69.02 ± 0.933 | D1: 98 ± 1.04  D2: 93 ± 1.04  D3: 98 ± 1.04  D4: 100 ± 1.04  D5: 100 ± 1.04  D6: 100 ± 1.04  D7: 98 ± 1.04  D8: 93 ± 1.04  D9: 95 ± 1.04  D10: 95 ± 1.04 |
| Karga & Mandal, 2016 | D1: 0.097 ± 0.004  D2: 0.051 ± 0.005  D3: 0.089 ± 0.003  D4: 0.092 ± 0.004 | D1: 22.62 ± 11.23  D2: 11.29 ± 4.82  D3: 20.41 ± 2.43  D4: 21.51 ± 4.17 |  | D1: 86.67 ± 1.67  D2: 78.33 ± 3.33  D3: 83.33 ± 4.41  D4: 81.67 ± 1.67 |
| Smith, 2013 |  |  | D1: 5.85 ± 1.1097  D2: 5.44 ± 0.82  D3: 5.33 ± 1.1671  D4: 4.19 ± 0.5782  D5: 4.53 ± 0.8266 |  |
| Lawrence, 2012 |  | D1: 723.09 ± 128.75  D2: 837.97 ± 32.67  D3: 900.69 ± 47.40  D4: 937.73 ± 79.38  D5: 221.46 ± 42.75 | D1: 97.92 ± 27.18  D2: 110.17 ± 18.42  D3: 112.05 ± 9.89  D4:  109.11 ± 8.67  D5: 57.29 ± 13.77 | D1: 96.9 ± 1.9  D2: 99.4 ± 0.6  D3: 98.8 ± 0.8  D4: 98.8 ± 1.6  D5: 98.8 ± 2.1 |

7.4. **Supplementary data 4**

The sub-group analysis findings for category 3 and 5 SGR are seen in Supplementary Data Fig. 1. Animal-based diets produced a significant effect on SGR between the studies (One-way ANOVA: F_(6, 360)_ = 492.6, *p* < 0.0001) (Supplementary Data Fig.1A). Lanes (2021) was found to produce the greatest increase in SGR compared to all other feeds with Tukey’s multiple comparison test. No difference was seen between Sevgili (2018) and Fernandes (2016) with 50% FM. Similarly, no significant difference was seen between Vural (2021) and Dhanasiri (2020) with FM. Comparatively, the student’s t-test determined no significant difference between 1.6% Royal Jelly compared to 6% FO and 0.5% CLEO in category 5 for SGR (student’s t-test: t(63) = 0.9485, *p* = 0.3465) ^26,36^ (Supplementary Data Fig. 1B).

**Supplementary Data Fig.1: The SGR sub-group analysis for categories 3 and 5.**

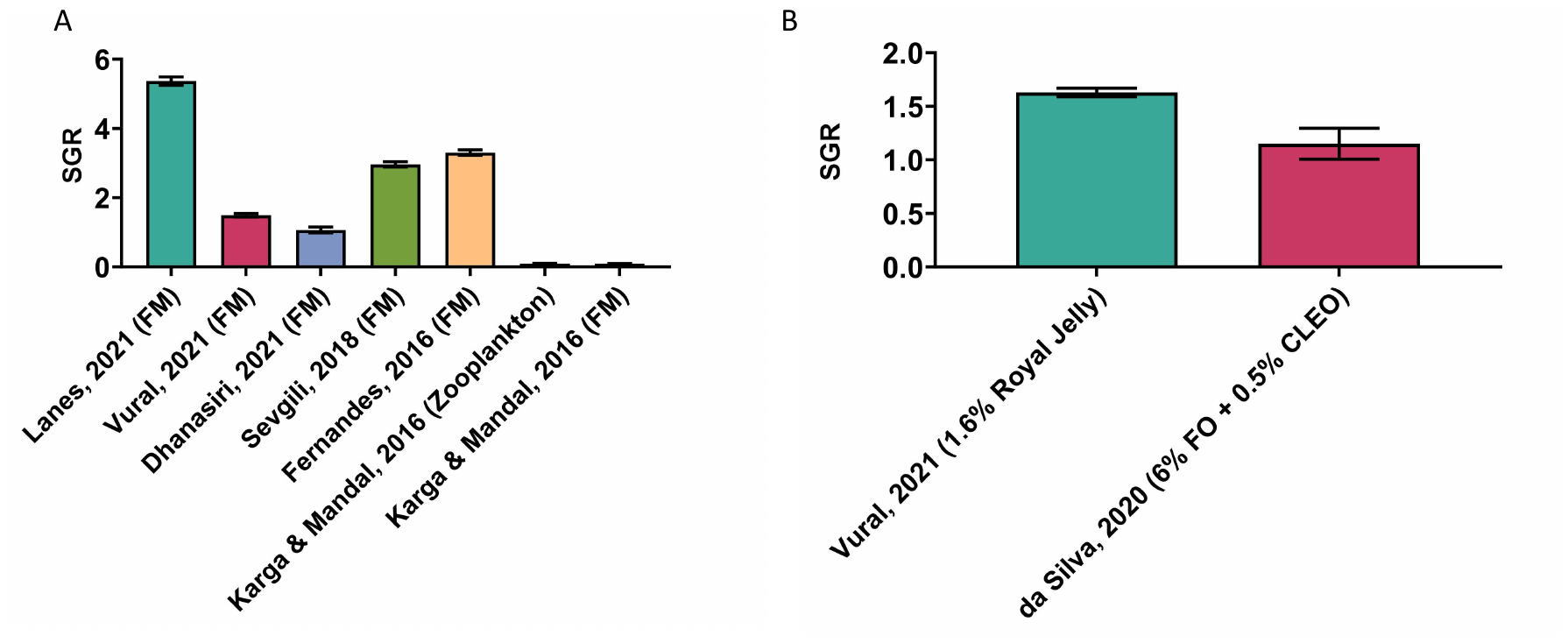

Note: The sub-group analyses for category 3 (A) and category 5 (B) SGR findings using a one-way ANOVA (A) and student’s t-test (B). The significance values are not included on the graphs due to too many significance values. Please see the GraphPad file on OSF for full analysis results.

7.5. **Supplementary data 5**

The sub-group analysis findings for categories 2-5 percentage weight gain are seen in Supplementary Data Fig. 2. A plant-based diet resulted in a significant effect on percentage weight gain with 100% Chlorella sp. causing a significant increase in weight gain compared to soy-protein (student’s t-test: t(83) = 15.44, p < 0.0001) (Supplementary Data Fig.2A). An animal-based diet also caused a significant effect on overall percentage weight gain (one-way ANOVA: F_(9, 630)_ = 24.28, *p* < 0.0001) (Supplementary Data Fig.2B). Tukey’s multiple comparison test confirmed Lanes (2021) to produce the greatest increase in weight compared to all other studies. Lanes (2021) was also found to produce the greatest weight gain in category 4 (one-way ANOVA: F_(2, 237)_ = 53.94, *p* < 0.0001) (Supplementary Data Fig.2C). No difference was seen between the two other insect-based diets. A significant effect on weight gain was seen in category 5 with 1.6% Royal Jelly significantly increasing percentage weight gain compared to a pro-biotic supplement (student’s t-test: t(13) = 24.09, *p* < 0.0001) (Supplementary Data Fig.2D).

**Supplementary Data Fig. 2: The sub-group analysis for categories 2-5 for weight gain (%).**

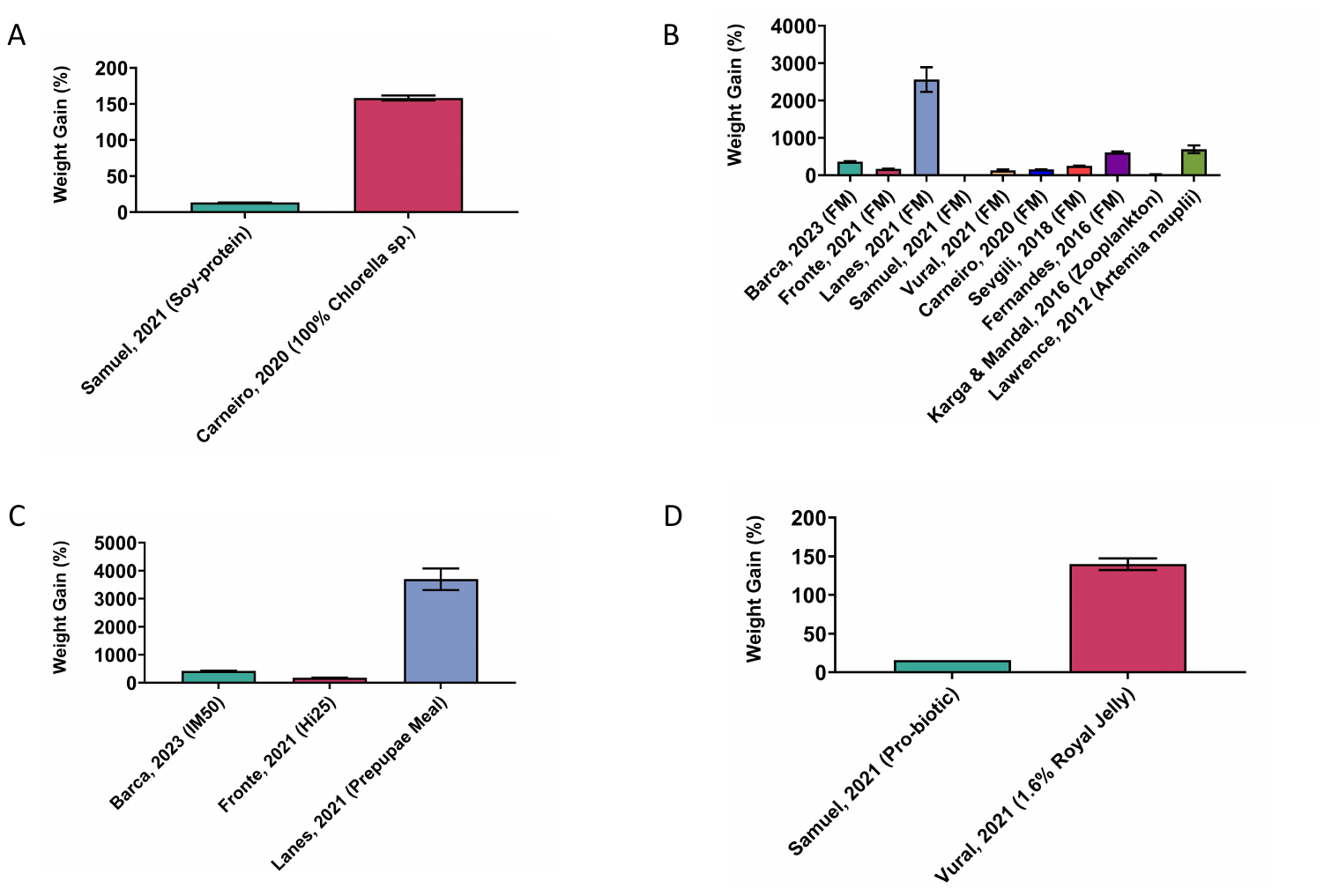

Note: The sub-group analyses for category 2 (A), 3 (B), 4 (C) and 5 (D) for weight gain (%) using a one-way ANOVA (B and C) and student’s t-test (A and D). The significance values are not included on the graphs due to too many significance values. Please see the GraphPad file on OSF for full analysis results.

7.6. **Supplementary data 6**

The sub-group analysis for category 3 and 6 for percentage length gain are seen in Supplementary Data Fig. 3. Lanes (2021) FM-based diet produced the greatest increase in percentage length gain (one-way ANOVA: F_(2, 167)_ = 62.73, *p* < 0.0001) (Supplementary Data Fig. 3A). The other FM-based diet also had significantly increased percentage length gain compared to the fish protein hydrolysate diet ^28,30^. A mix of protein sources also significantly impacted length gain results (student’s t-test: t(620) = 10.83, *p* < 0.0001) (Supplementary Data Fig. 3B).

**Supplementary Data Fig. 3: The sub-group analysis for categories 3 and 6 for length gain (%).**

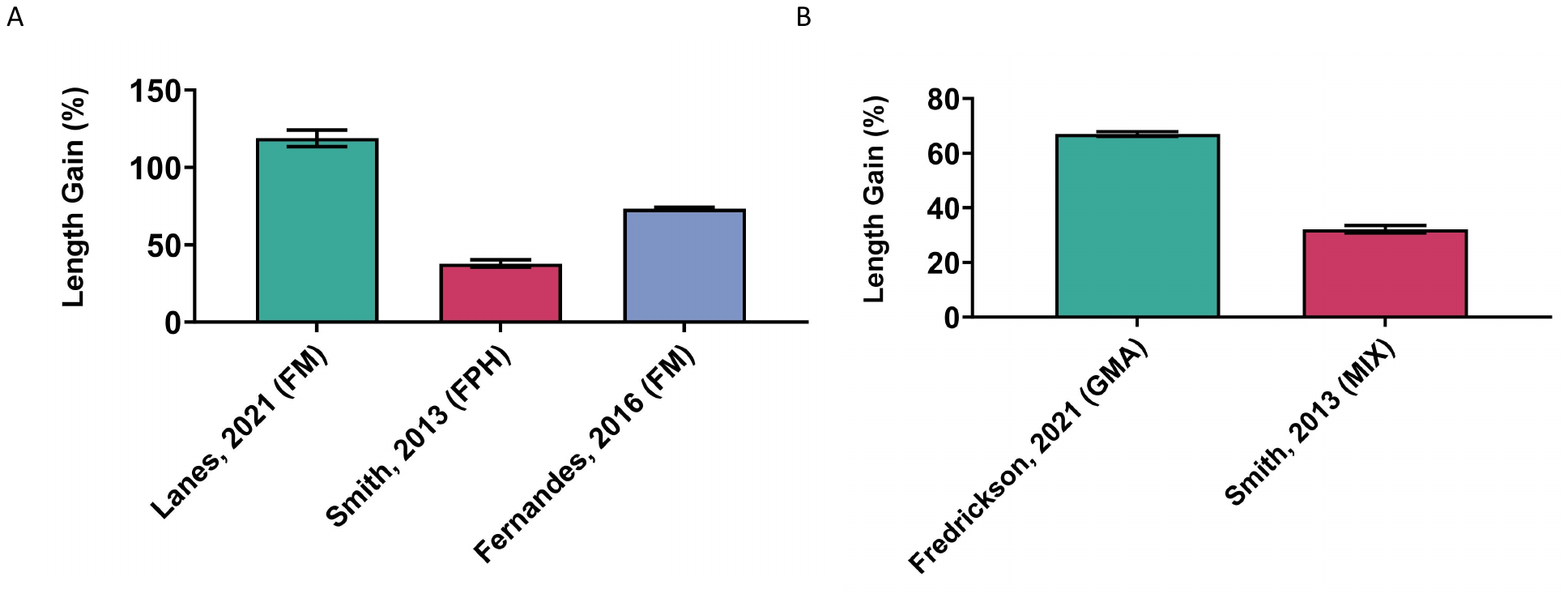

Note: The sub-group analyses for category 3 (A) and 6 (B) for length gain (%) using a one-way ANOVA (A) and student’s t-test (B). The significance values are not included on the graphs due to too many significance values. Please see the GraphPad file on OSF for full analysis results.
